# Patient-informed biomechanical modelling reveals mechanical mechanism of brain damage in idiopathic normal pressure hydrocephalus

**DOI:** 10.64898/2026.04.27.721128

**Authors:** Vahid Darvishi, Martina Del Giovane, Michael C. B. David, Magdalena A. Kolanko, Anastasia Gontsarova, Paresh A. Malhotra, Christopher Carswell, David J Sharp, Mazdak Ghajari

## Abstract

Idiopathic normal pressure hydrocephalus (iNPH) is a globally growing neurological disorder in older adults, radiologically characterised by enlargement of ventricles. However, it remains unknown whether ventricular enlargement can produce biomechanical loading large enough to drive brain morphological changes and tissue damage. Here, we develop an anatomically detailed biomechanical model of ageing human brain and apply ventricular enlargement using a three-dimensional displacement field derived from MRI of iNPH patients and age-matched controls. The model accurately reproduces radiological markers of iNPH, including Evans index, callosal angle and high-convexity sulcal narrowing. It further predicts large mechanical strains in periventricular white matter, particularly within the corpus callosum and anterior thalamic radiations, tracts consistently implicated in iNPH imaging abnormalities. These findings provide strong evidence that ventricular enlargement induces mechanical strain that contributes to iNPH brain abnormalities, which can potentially be reversed by reducing strain following shunting surgery. The biomechanical brain model forms the foundation of a predictive digital platform and future “digital twin” technology to support diagnosis, patient stratification and treatment planning in iNPH.

**Key Points:**

- We develop an anatomically detailed biomechanical model of the brain, incorporating sulci, septum pellucidum and all four ventricles.
- A novel data-driven loading approach is introduced which uses 3D displacement fields from finite element-based registration of healthy and iNPH patient MRI, replacing the arbitrary pressure gradients of previous models.
- The model predictions closely match established radiological markers measured in iNPH patients, including Evans index, callosal angle and high-convexity sulcal narrowing.
- Ventricular enlargement generates large mechanical strains concentrated in periventricular white matter, providing a biomechanical explanation for the structural abnormalities observed in iNPH.

## Introduction

Idiopathic normal pressure hydrocephalus (iNPH), first introduced by Hakim and Adams in 1964, is a neurological disorder characterized by abnormal enlargement of the ventricles and symptoms including gait disturbance, urinary incontinence, and cognitive impairment, also known as Hakim’s triad ^1–6^. iNPH primarily affects individuals over age 60, with a prevalence of 1.5–2.5% in those over 65 and up to 6% in over 80 ^7,8^. With the global population aged 65 and older projected to double by 2050, the burden of iNPH will likely increase significantly in the coming decades ^9^.

iNPH is often misdiagnosed as Alzheimer’s disease (AD) due to their overlapping symptoms and radiological features ^10,11^. However, unlike AD, iNPH can be reversed in some cases ^12^. Cerebrospinal fluid (CSF) shunting improves symptoms ^13,14^, and approximately 15% of all shunt procedures in the UK are performed for iNPH ^15^. Yet, the reasons as to why a proportion of patients do not benefit from shunting remain unclear, likely due to limited understanding of the disease’s underlying mechanisms ^3,4^.

iNPH is likely to have a biomechanical nature, distinguishing it from hydrocephalus ex vacuo, where ventricular enlargement occurs passively to fill the space left by shrinking brain tissue, as commonly seen in AD. Studies have suggested that CSF accumulation within the ventricles applies pressure on the ventricular walls, contributing to their expansion and brain deformation ^16–20^. In addition, early changes in the mechanical properties of brain tissue leading to secondary ventricular enlargement has also been suggested as another explanation for iNPH ^21^. However, limitations in experimental and clinical methods for measuring biomechanical variables such as pressure gradients and brain deformation make it difficult to investigate potential biomechanical mechanisms involved in iNPH ^22^.

While the mechanisms underlying iNPH remain unclear, their effects consistently manifest as distinct macroscopic radiological features that form the basis of clinical diagnosis of iNPH. Ventricular enlargement, the principal radiological sign of iNPH, is typically quantified using the Evans index (EI) and z-Evans index (z-EI), which measure the height and width of the ventricles, and callosal angle, which assesses the angle between the two lateral ventricles ^23,24^. A narrowing of the CSF space at the high midline and high convexity areas are other radiological characteristics of the disease ^19,25,26^. Although these features are routinely used in clinical practice, it remains unclear how ventricular enlargement leads to such characteristic morphologies and whether these macroscopic changes are associated with tissue damage, particularly white matter abnormalities.

Several studies have identified white matter abnormalities in periventricular regions in iNPH patients by using diffusion weighted imaging (DWI) ^27–32^. Axonal injury is thought to underlie the DWI abnormalities observed in white matter ^4^, representing microscopic hallmarks of iNPH. It has also been suggested that tracts located in periventricular areas are related to the clinical symptoms of iNPH. For instance, the corpus callosum, anterior thalamic radiations and corticospinal tract have been associated with gait disturbance ^27,33–36^, and the anterior thalamic radiations and cingulate cortex and its related tracts have been linked to urinary incontinence ^37^. The spatial proximity of these tracts to regions of ventricular expansion suggests that mechanical factors may contribute to their abnormalities. In parallel, mechanical strain, which is a measure of local brain tissue deformation, has been used as a marker of tissue damage in traumatic brain injury, where regional strain distribution has been shown to predict the location of brain tissue damage and explain symptoms ^38–41^. These findings support the possibility that mechanical loading may contribute to the white matter structural abnormality and symptoms of iNPH.

Computational models that incorporate anatomy and biomechanical properties of different tissues hold promise in investigating the relationship between brain mechanical loading and structural abnormalities in iNPH. Biomechanical computational models of iNPH have been developed in recent years. While some models incorporate a realistic description of brain anatomy ^42–46^, most focus on predicting CSF velocity or pressure field, rather than brain tissue deformation. A few studies focused on predicting brain deformation and tissue damage indicators, such as strain distribution, but their models lack key anatomical features ^44,46– 49^. For instance, none of the previous models have incorporated 3^rd^ and 4^th^ ventricles and their expansion during iNPH. They also have not incorporated the entire anatomy of lateral ventricles, particularly the inferior regions. Moreover, they have used MRI scans of young adults, thus not accounting for the differences between younger and older healthy brains. Another limitation of previous computational models of iNPH is the loading condition used to simulate ventricular expansion. They expand the ventricles by applying an inaccurate or arbitrary transmantle pressure gradient (the pressure difference between the cranial subarachnoid space and the lateral ventricles) on the boundary of ventricles. However, this approach produces an unrealistic shape of expanded ventricles, leading to an inaccurate prediction of brain deformation ^44^. As such, no computational models have been able to predict the key radiological features of iNPH, nor have they explained the characteristic white matter abnormalities seen in iNPH.

Here we develop a new patient-informed computational model of iNPH, which incorporates fine anatomical details of key brain regions, including all four ventricles and sulci. We also introduce a new loading condition derived from MRI scans of iNPH patients and age-matched healthy controls to accurately capture the morphology of enlarged ventricles. We evaluate whether this novel biomechanical model can accurately predict key radiological markers of iNPH. In addition, motivated by the spatial correspondence between ventricular morphology and periventricular white matter abnormalities in iNPH patients, we investigate the validity of the model’s mechanical strain predictions in identifying the white matter tracts often reported as abnormal in previous studies ^50,51^. This study can improve our understanding of the role of biomechanics in iNPH and establish the computational modelling framework as the foundation for a digital platform for improving iNPH diagnosis, patient stratification and planning of shunting interventions.

## Results

### The anatomically detailed finite element model of brain

The MRI scans of 30 healthy controls (HC) (16 males, mean age = 73.80±5.71) were averaged to create a template. An in-house pipeline was then used to convert the template into a finite element (FE) model of the brain ^41^. Anatomical features, including sulci, pia, falx and tentorium, were also generated using the pipeline (Fig. 1a and Fig. 1b). Manual segmentation of the ventricles and septum pellucidum (SP) of the template resulted in an accurate representation of their fine anatomy (Fig. 1c and Fig. 1d). This included the lower parts of the ventricles, which are missing in previous models ^44,46–49^. Hyperelastic isotropic Ogden material models with constants extracted from experiments on human brain tissues were assigned to different anatomical regions to predict nonlinear mechanical response of the tissues to loading ^52,53^.

**Fig. 1.**
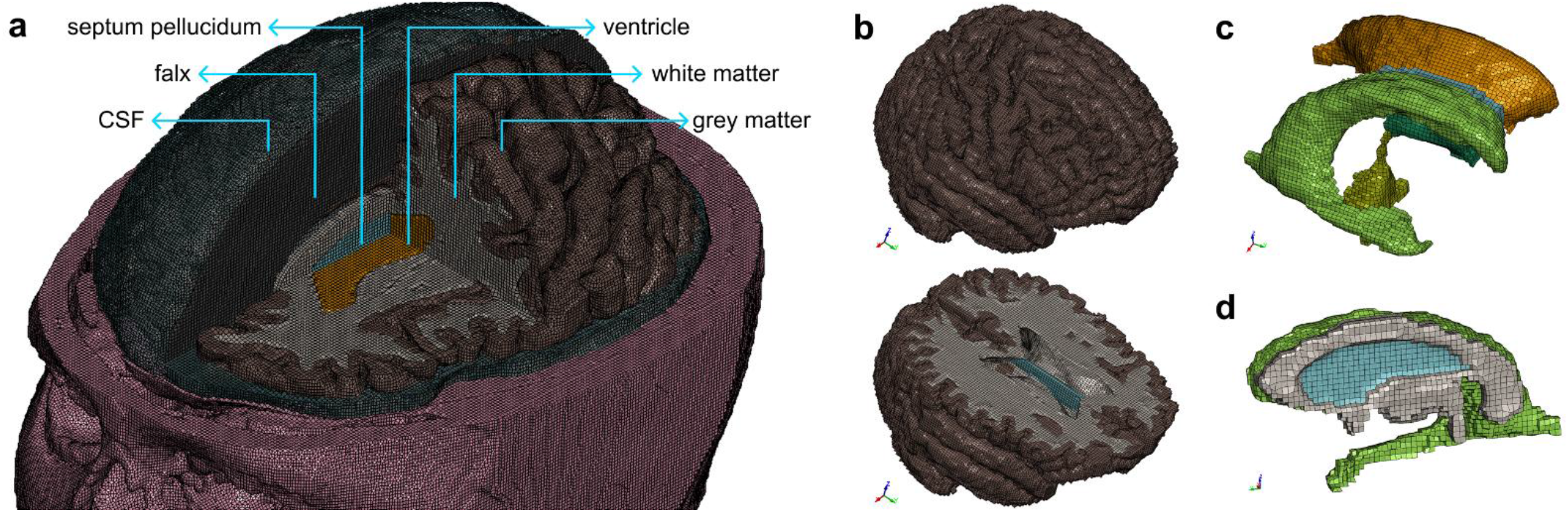
The anatomically accurate finite element model of the brain, representing healthy old adults. **a**, Geometry of key brain regions after incorporation of anatomical features, including sulci, falx, septum pellucidum (SP), and pia mater, **b**, Detailed representation of gyri and sulci, **c**, Refined and anatomically accurate ventricular system used in this study and **d**, Anatomy of SP shown in blue.

### Patient-informed ventricular expansion

To reproduce ventricular expansion, a three-dimensional displacement field was applied to the lining of the ventricles. This displacement field was obtained from mapping the ventricles of the HC template to the ventricles of an iNPH template, created from the MRI scans of 22 iNPH patients (14 males, mean age = 71.91±7.06). Compared with linear and nonlinear image registration, the FE-based registration provided superior alignment with iNPH ventricles (Fig. 2). Using the node-wise surface comparison, we found a 95^th^ percentile Hausdorff distance (HD95) ^54^ of 2.2 mm for the FE-based registration, versus 12.3 mm for the linear and 4.7 mm for the nonlinear image registrations. Hence, the FE-based registration was used to determine the displacement field of ventricular linings.

**Fig. 2.**
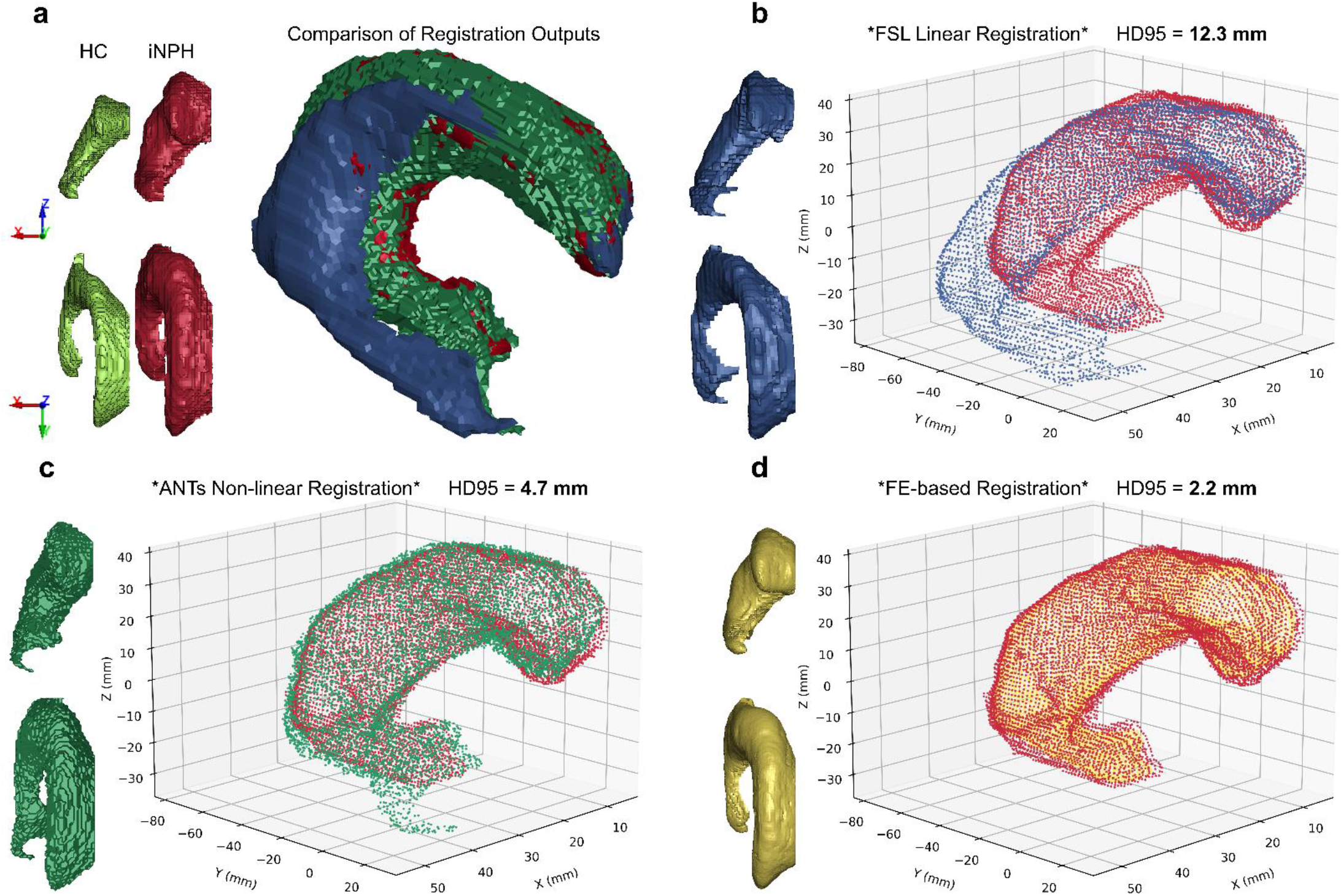
Different registration methods for aligning HC and iNPH ventricles. This figure shows the registration and shape analysis of the right lateral ventricle. **a**, Anterior-posterior and superior-inferior view of right ventricle of HC (green) and iNPH (red), along with an isometric view of the registered versions of the right ventricle. **b**, FSL linear registration result (blue), evaluated against iNPH. **c**, ANTs nonlinear registration output (dark green), evaluated against iNPH. **d**, FE-based registration output (yellow), evaluated against iNPH.

The lining of ventricles was made of approximately 21,000 finite element nodes. The displacement fields were converted into displacement vectors, which were used to prescribe the displacement of these nodes. Ventricular expansion was modelled by increasing the magnitude of these vectors simultaneously from zero to their maximum value (Fig. 3).

**Fig. 3.**
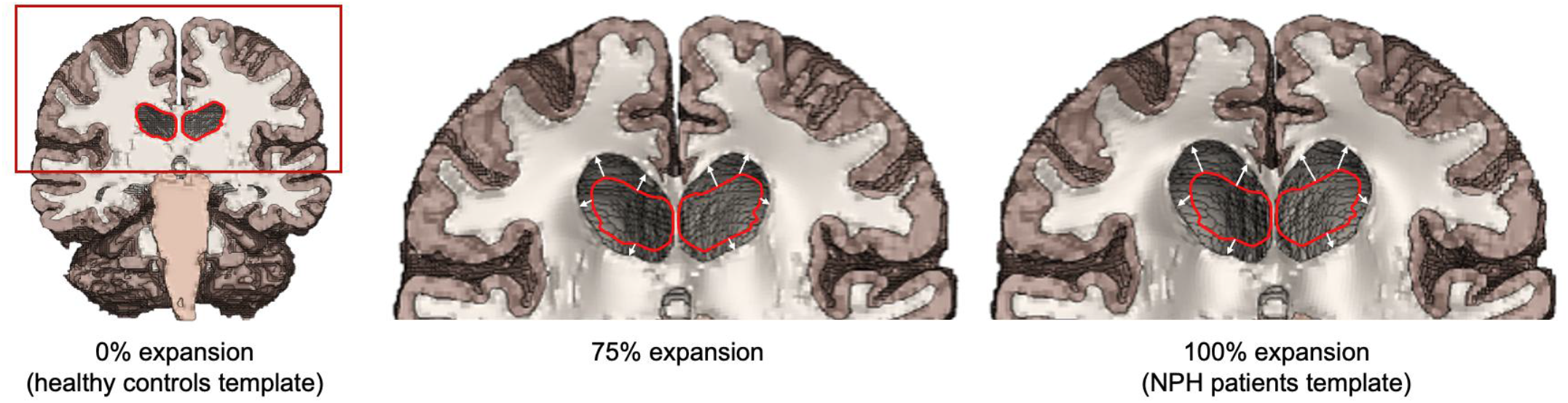
Simulation of ventricular expansion by using a displacement field. The red outlines show the lining of ventricles in the HC template. The white arrows show the displacement vectors, determined from the FE-based registration of the ventricular lining of HC to iNPH.

### The model reproduces ventricular morphology metrics

Evans index (EI) and z-Evans index (z-EI) are widely used radiological markers for iNPH. EI is defined as the ratio of the maximum distance between the frontal horns of the lateral ventricles to the maximum internal diameter of the cranium in the lateral direction (Fig. 4a). Similarly, the z-EI is defined as the ratio of the maximum z-axial distance between the upper and lower parts of the frontal horn to the maximum internal diameter of the cranium in the z direction, measured at the posterior margin of the interventricular foramen (Fig. 4b). These indices were measured in iNPH patients by a neuroradiologist and also predicted by the brain model during ventricular expansion simulation (Fig. 4c and Fig. 4d). The EI of the FE model increased linearly from 0.283 to 0.345 as the ventricles fully expanded, falling within the range measured in patients and differing by only 4.4% from the patients’ median (Fig. 4e). The z-EI also increased linearly from 0.306 to 0.417, with a deviation of just 1.8% from the patients’ median (Fig. 4f).

**Fig. 4.**
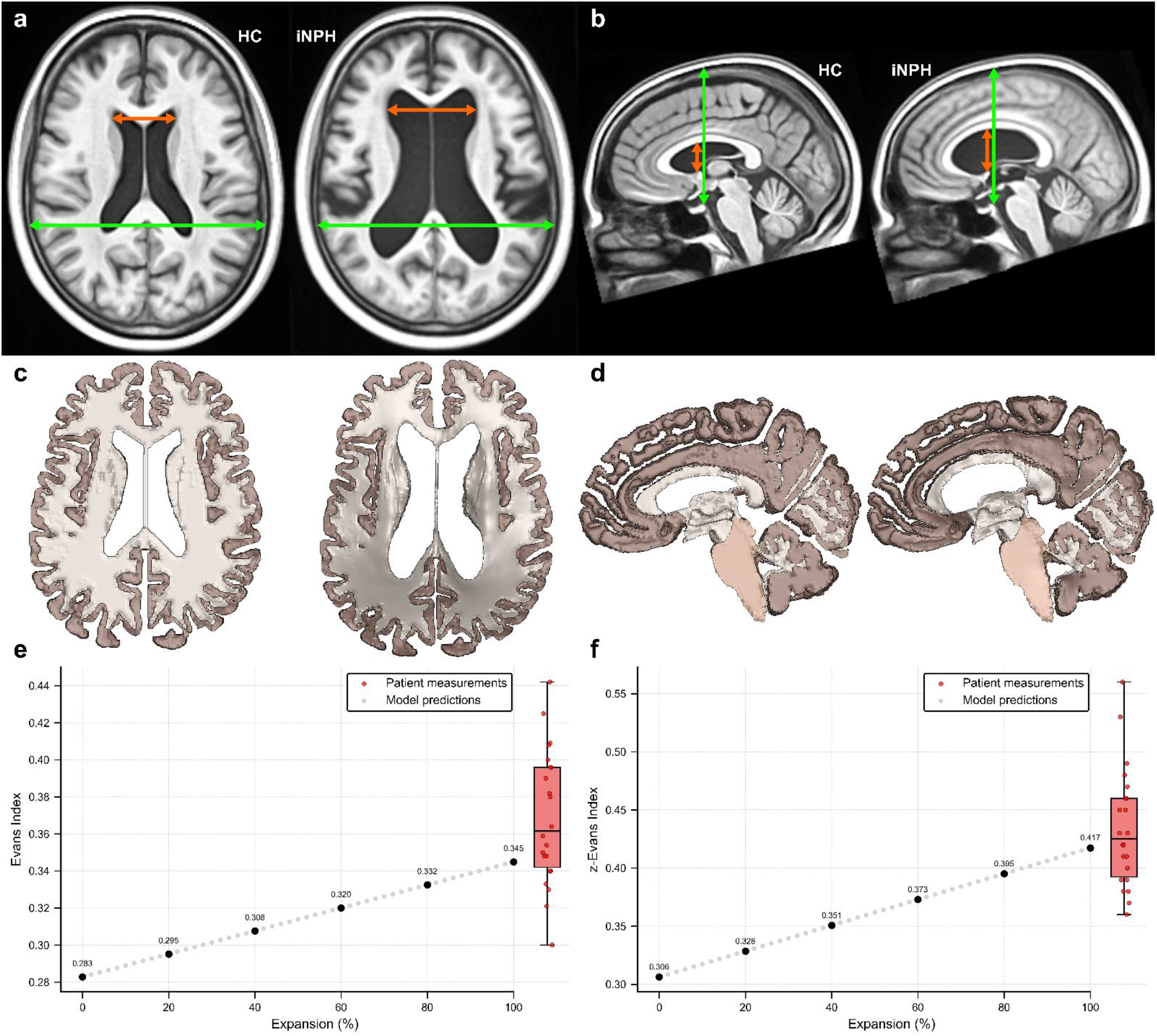
Comparison of EI and z-EI between MRI templates and the brain model. **a**, Definition of EI. **b**, Definition of z-EI. **c**, Increase in the EI predicted by the model. **d**, Increase in the z-EI predicted by the model. **e**, EI measurements show a close agreement between data extracted from iNPH patients’ scans and model predictions. **f**, z-EI measurements show a close agreement between data extracted from iNPH patients’ scans and model predictions.

The reduction in the callosal angle, the angle between the lateral ventricles, is another indicator of iNPH (Fig. 5a) ^55–57^. This angle is measured on the coronal plane perpendicular to anterior commissure - posterior commissure line at the level of the posterior commissure. The FE model predicted a slightly nonlinear reduction in the callosal angle, decreasing from 116◦ before ventricular expansion to 71.5◦ at full expansion, corresponding to only a 0.7% relative error compared with the median callosal angle of iNPH patients (Fig. 5b and c).

**Fig. 5.**
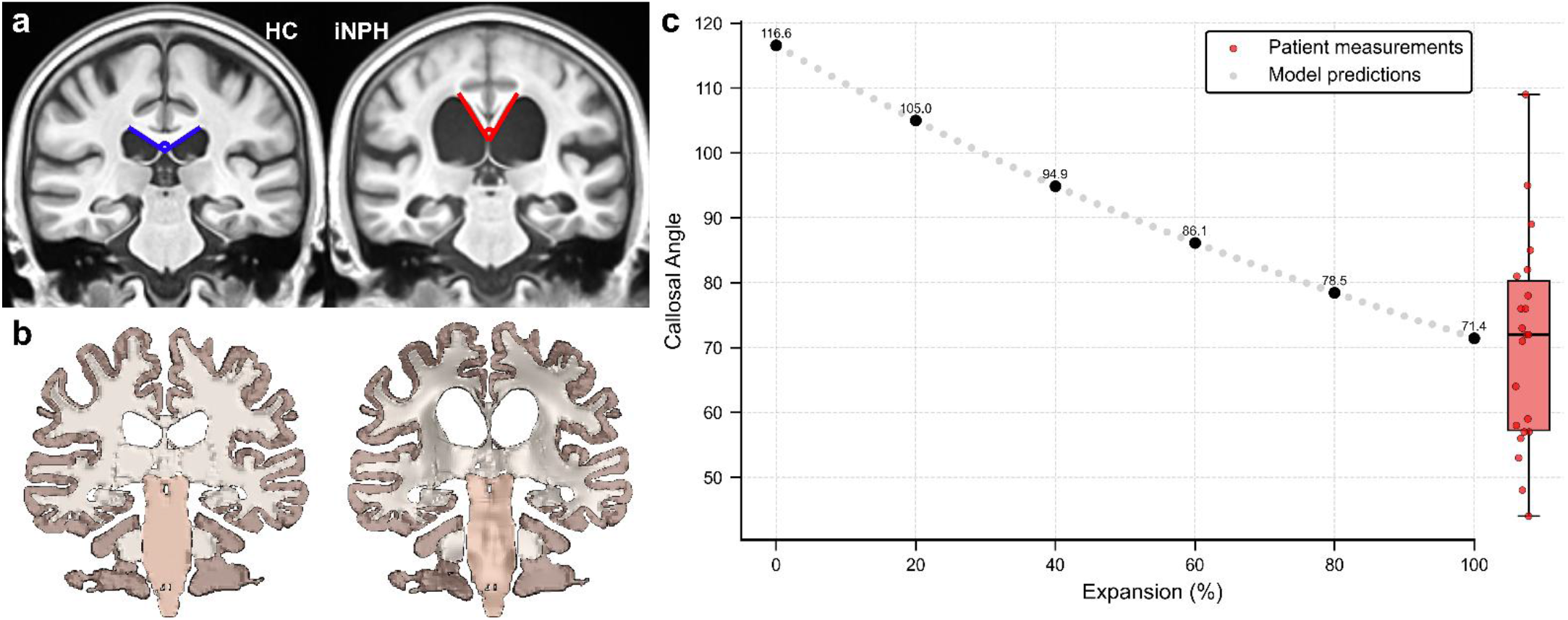
Callosal angle during expansion of ventricles in iNPH. **a**, Definition of Callosal angle and its measurement in MRI templates. **b**, Reduction of callosal angle predicted by the model. **c**, Change in callosal angle over time during ventricular expansion predicted by the model which is in a close agreement with measurements from iNPH MRI scans.

### Narrowing of interhemispheric fissure and sulci

We also examined broader brain morphological changes induced by ventricular expansion. A reduction in the CSF space at high midline and high convexity areas have been reported in iNPH patients, seen on MRI as narrowing of the interhemispheric fissure and superior cortical sulci ^19,25,26^. Consistent with these observations, the brain model predicts a progressive decrease in the distance between the left and right hemispheres as the ventricles expand, with the greatest reduction in the posterior regions (Fig. 6a). Likewise, the model predicts narrowing of sulci in high convexity areas, which again is more pronounced in the posterior regions (Fig. 6b).

**Fig. 6.**
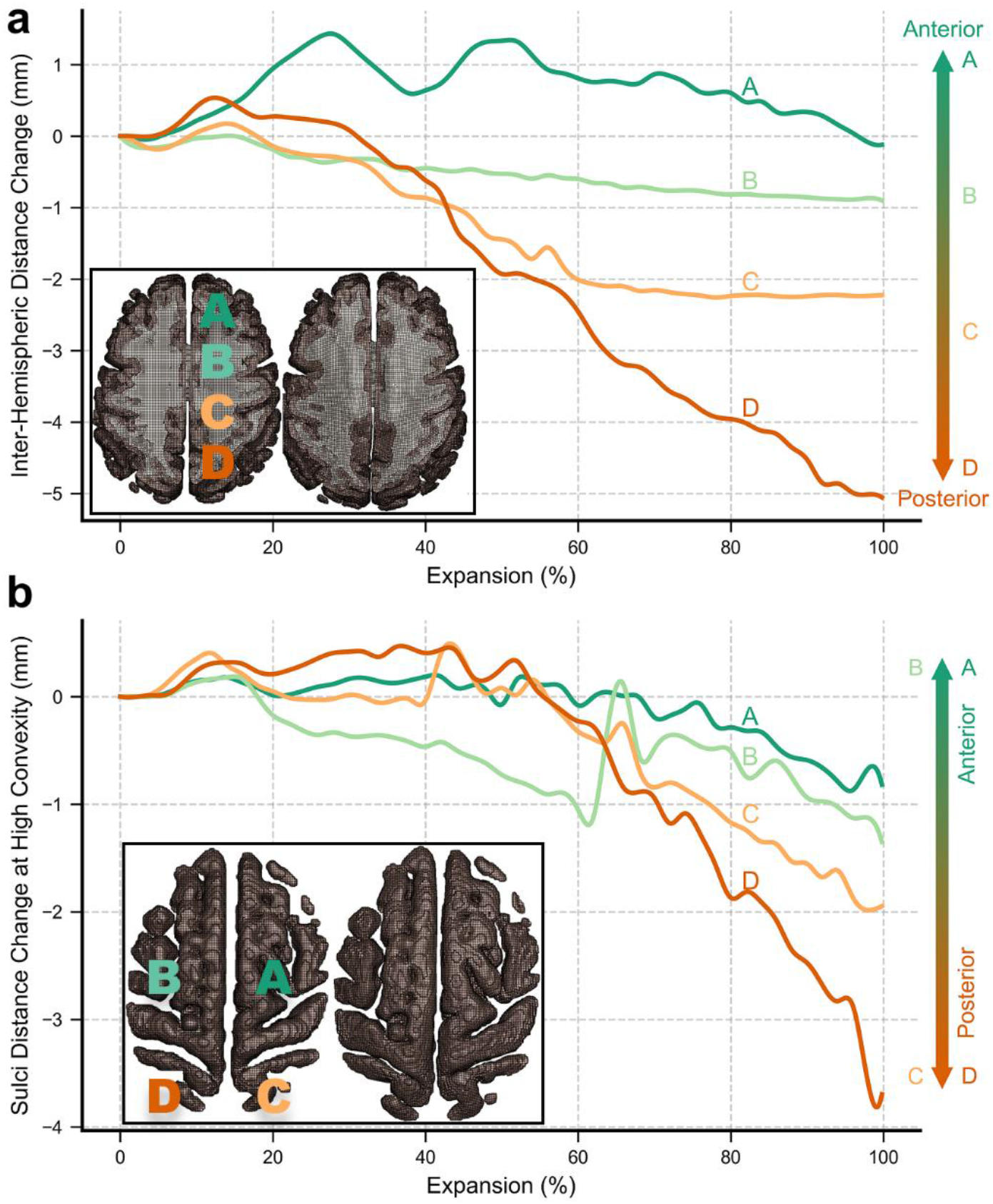
Predicting interhemispheric fissure and sulci narrowing at high convexity in iNPH. **a**, Interhemispheric distance reduction at multiple locations during the ventricular expansion, with the greatest reduction in posterior locations. **b**, Narrowing of the sulci at high convexity, which is more pronounced in posterior regions.

### Reduction in septum pellucidum thickness

The septum pellucidum (SP), the thin layer between the lateral ventricles, has been highlighted as a potential region of interest for brain-related neurological conditions ^58^. We observed a reduction in SP thickness when comparing the HC and iNPH MRI templates, with the reduction being more pronounced in the posterior region (Fig. 7a). The brain model predicted that the SP thickness decreases nonlinearly as the ventricles expand, with greater thickness reduction in the posterior region of the SP (Fig. 7b).

**Fig. 7.**
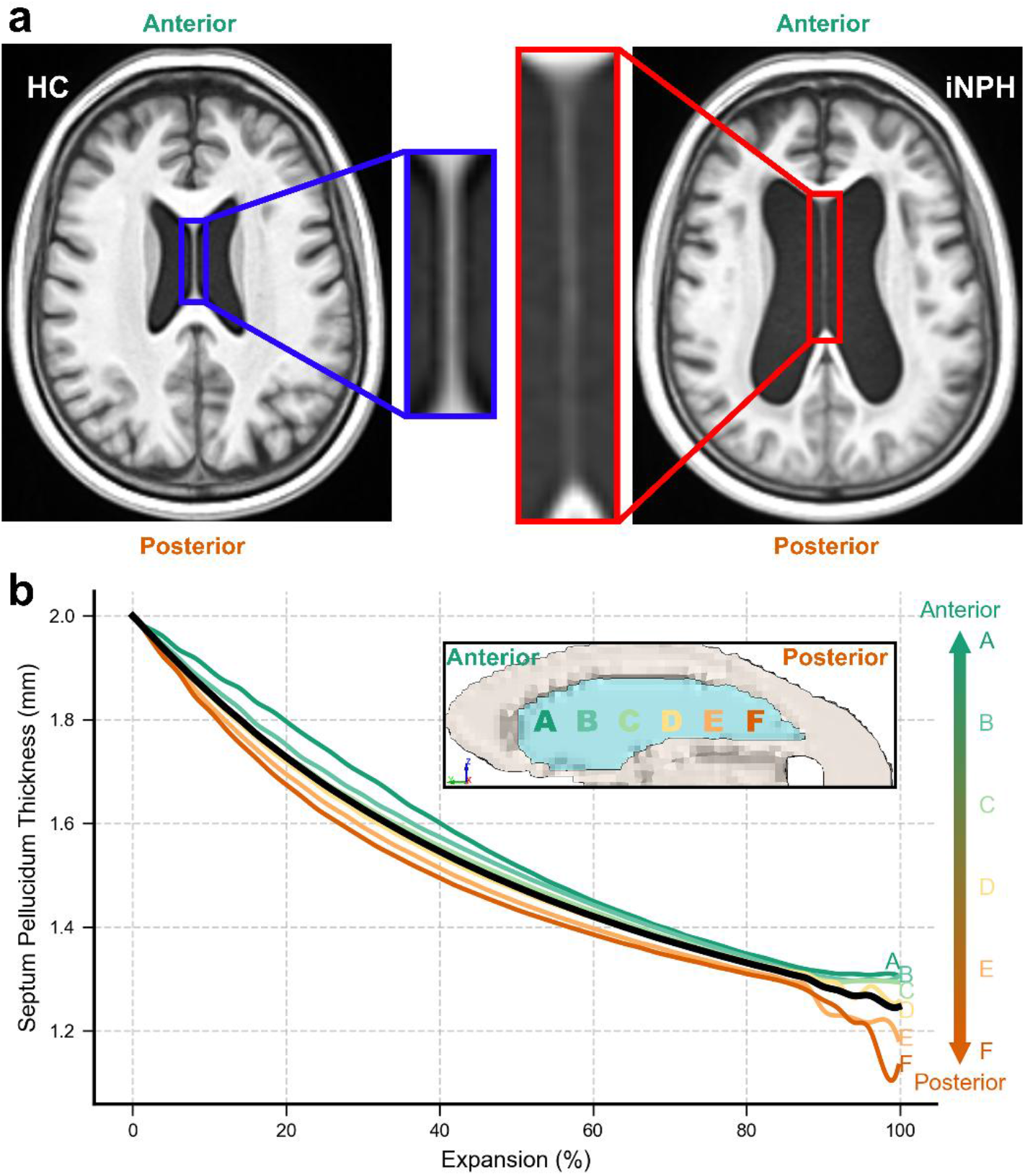
SP thickness reduction during ventricular expansion observed in MRI scans of patients and predicted by the brain model. **a**, SP thickness in HC and iNPH MRI templates, showing reduction in iNPH SP thickness, which becomes more pronounced in the posterior regions. **b**, The brain model predicts a continual reduction in SP thickness during iNPH progression, with reductions more pronounced in posterior regions. The black line represents the average reduction.

### Large strains in areas of white matter abnormality

The distribution of maximum principal Green-Lagrange strain across the brain, produced by ventricular expansion, was predicted by the brain model. The largest strains were concentrated in the periventricular regions and most notably in the septum pellucidum and corpus callosum (Fig. 8a). To determine strain distributions in white matter tracts, the JHU ICBM-DTI-81 white matter tract atlas was used ^59^. The 95^th^ percentile strain in white matter tracts ranged from 0.09 to 0.64 (Fig. 8b). The tracts with largest 95^th^ percentile strain were corpus callosum genu, splenium and body, and anterior thalamic radiations.

**Fig. 8.**
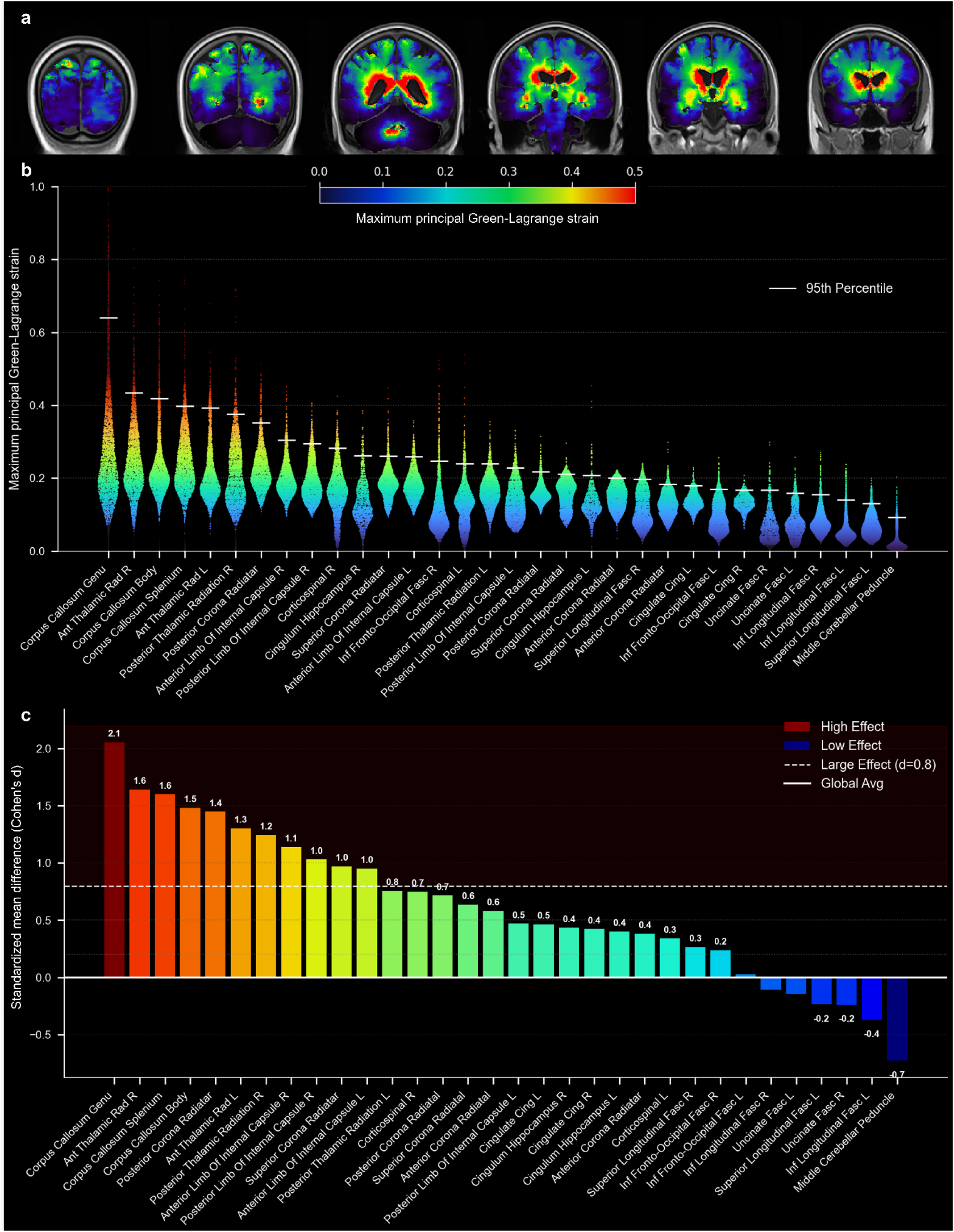
Maximum principal Green-Lagrange strain distribution predicted by the iNPH brain model. **a**, Maximum strain in different regions of the brain. **b**, Strain distribution across white matter tracts, based on maximum principal strain analysis. **c**, Standardized mean difference comparison between tracts, showing the corpus callosum and anterior thalamic radiations experience significantly higher strain compared to other tracts.

To identify the white matter tracts experiencing significantly higher levels of strain compared to the whole brain average, the effect size (Cohen’s d) was calculated. The Cohen’s d is a measurement of the magnitude of the difference between the tract’s strain and the global average, expressed relative to the overall variability in strain. Largest effect sizes were measured for the corpus callosum genu, splenium, and body, and the anterior thalamic radiations, with Cohen’s d values of 2.1, 1.6, 1.5, and 1.6 respectively (Fig. 8c).

## Discussion

We show that an anatomically detailed biomechanical model of brain, loaded by displacement of the ventricular lining, can accurately reproduce established radiological markers of the iNPH disease, including Evans index, z-Evans index and callosal angle. Moreover, the model successfully predicts morphological changes seen in MRI of iNPH patients, including narrowing of high convexity sulci and interhemispheric distance reduction. In addition, the model predicts large mechanical strains in white matter tracts where abnormalities have been seen in diffusion imaging of iNPH patients. Together, these findings indicate that brain biomechanical loading resulting from enlargement of ventricles plays an important role in iNPH pathophysiology. They also validate model predictions, supporting the use of this model in future studies on understanding iNPH mechanisms and improving its diagnosis and treatment.

A significant advancement of the presented brain model, compared with previous models of iNPH ^43–46^, is incorporating anatomical features that influence brain deformation during ventricular enlargement, such as sulci and falx. The falx may have a role in constraining adjacent areas such as the corpus callosum during ventricular expansion. In contrast, sulci and CSF space at high convexity areas provide space for brain tissue movement as ventricles enlarge, thus reducing brain deformation and strain in these areas. In addition, an accurate definition of ventricular anatomy, including the lower parts of the lateral ventricles and 3^rd^ and 4^th^ ventricles, was incorporated in the model. This enabled the model to predict areas of elevated strain in the vicinity of ventricles as they enlarge (Fig. 8a). This accurate and detailed prediction of neuroanatomical distribution of mechanical strain may help understand abnormalities observed in iNPH.

Another key novelty of the modelling approach is using MRI data to displace the ventricular lining, thus allowing for an accurate reproduction of the morphology of expanded ventricles. Previous models have failed to predict the shape of the expanded iNPH ventricles because they used pressure gradients to derive ventricular expansion, leading to a uniform expansion across ventricular regions ^43–46,48,49^. The registration of HC to iNPH ventricles provided around 21,000 unique displacement vectors for nodes of ventricles’ linings. This method allowed for the accurate prediction of nonuniform expansion of the ventricles, closely reproducing the heterogeneous ventricular enlargement seen in iNPH patients, with excellent predicted values for the established radiological metrics of iNPH. The model demonstrated higher predictive accuracy for the z-EI and callosal angle compared to the EI, which is due to the sensitivity of these measurements to initial segmentation boundaries and the surface smoothing required for numerical stability in the FE models.

The model predicted additional morphological changes, including narrowing of sulci at the high convexity, narrowing of the interhemispheric fissure and thinning of the septum pellucidum (SP). These changes were most pronounced in the posterior regions. Tight high convexities and compressed midline subarachnoid spaces are morphological features that suggest iNPH ^19,25,26^. The model prediction of these features further supports the role of mechanical loading in driving iNPH morphological changes. The thinning of the SP has not been widely reported as an imaging feature of iNPH, possibly due to its small size, which complicates its measurement on MRI. Nonetheless, SP has been implicated in other neurological disorders ^58^. The model predicts a substantial average reduction of approximately 37% in SP thickness due to ventricular enlargement, suggesting that SP thickness may be a useful morphological feature in the iNPH context.

The model predicted large mechanical strains in periventricular white matter tracts, with the corpus callosum and anterior thalamic radiations sustaining the highest 95^th^ percentile strain values. This observation was further supported by the effect size analysis, which identified these tracts as experiencing significantly higher strain levels than the global brain average (Cohen’s d > 1.5). This aligns with MRI studies of iNPH patients, which consistently report diffusion abnormalities in periventricular regions, with the corpus callosum being reported in several studies ^27,28,31,60^. For example, one study comparing diffusion imaging of iNPH and Alzheimer’s disease patients reported significant reduction in fractional anisotropy, which indicates abnormality, in the corpus callosum ^25^. Consistent with these observations, we predicted highest levels of strain in corpus callosum. The relationship between mechanical stretching of white matter tracts and subsequent axonal injury has been established in animal studies ^39,61,62^. Our results further confirms the ability of the computational model in predicting brain abnormalities in iNPH.

The biomechanical model predicted highest strains in the corpus callosum and anterior thalamic radiations. These tracts have been most consistently associated with gait disturbance, an early and common symptom in iNPH ^33,63,64^. This unique pattern of strain distribution may provide a biomechanical explanation for the clinical progression of iNPH symptoms. It suggests that the early appearance of gait disturbance could be a consequence of early and continued mechanical stretching of tracts associated with gait. Animal studies have shown that stretching white matter tracts beyond a threshold can initiate functional deficits, and continued deformation can result in irreversible axonal damage ^62,65,66^. Our model predicts monotonically increasing strains in all tracts as ventricles expand, meaning that eventually more tracts can reach a threshold of irreversible damage. This raises the possibility that interventions aimed at reducing ventricular expansion may not be effective if tracts are permanently damaged, a potential reason for not responding to shunting in some patients.

This study highlights the key role of biomechanics in iNPH. While the brain model is purely biomechanical and does not account for potential biological or genetic factors, such as pre-existing neurodegeneration or tissue softening prior to ventricular enlargement, the strong agreement between model predictions and common morphological and structural indicators of iNPH suggests that mechanical forces play an important role in deriving the observed abnormalities. It also implies that ventricular enlargement imposes mechanical loading on brain tissue, rather than being solely a consequence of early tissue softening. Clinical interventions that reduce CSF volume and pressure, such as shunting procedures or CSF-suppressing pharmacologic treatments, have been shown to alleviate both imaging markers of white matter damage and patient symptoms ^67^. This therapeutic effect further supports the role of biomechanics in iNPH, where excessive CSF accumulation increases pressure at the ventricular surface, leading to tissue deformation and damage, which can be partially reversed when the mechanical loading is reduced ^31^.

We used MRI templates of healthy controls and iNPH patients to develop the brain model and prescribe ventricular enlargement for the model. While the model was able to predict common iNPH morphological changes and abnormal tracts, it fails to capture differences between patients. For instance, although the reduction in interhemispheric distance and narrowing of sulci at the high convexity have been reported in the literature as a common feature of iNPH ^19,25,26^, these features may not exist in MRI scans of all iNPH patients.

This can be related to the variations in ventricular enlargement or brain anatomy across patients, which can be tested by developing patient-specific models. These models should incorporate individualised neuroanatomy and ventricular enlargement description. A barrier against developing patient-specific models is the highly time-consuming FE-based registration, which was used to determine the displacement field of ventricular linings, achieving a successful registration with HD95 of 2.2 mm. A promising replacement for this approach is machine learning, which can reduce the simulation time by orders of magnitude without compromising the accuracy ^68,69^. This will bring the digital model described in this study a step closer to a digital twin of an iNPH patient, which can help with patient stratification and informing shunting procedure.

In conclusion, this study introduces a novel finite element model of iNPH that leverages detailed brain anatomy and a displacement-based loading approach to reproduce ventricular expansion with high accuracy. The model is validated by accurate prediction of key morphological changes and tissue abnormalities observed in iNPH, supporting the role of mechanical loading in iNPH pathophysiology. This computational model can be used to investigate the mechanical effects of ventricular enlargement on white matter tracts. Future studies should apply the model to the brain of NPH patients who underwent CSF drainage to assess the reversibility of strain caused by hydrocephalus and the associated brain abnormalities. This holds promise in patient stratification and for informing shunting procedure.

## Methods

### Brain MRI templates

Standard brain templates, which are average MRI images of healthy individuals, are commonly used for computational model development ^70^. However, these templates are for healthy young adults, whereas iNPH predominantly affects elderly individuals aged 60 and above. To represent the brain anatomy of healthy elderly adults, we created a template by averaging T1-weighted MRI scans of 30 healthy individuals aged 62–82 years (16 males, mean age = 73.80±5.71) using antsMultivariateTemplateConstruction.sh from the ANTs library (Fig. 9a) ^71^. An initial average template was generated by registering all subjects to one reference subject, and this average was then registered to the MNI152 atlas affinely. All scans were subsequently realigned to the evolving template using rigid registration with three iterations to improve accuracy. This process was then repeated with affine and finally symmetric diffeomorphic accordingly to further refine the averaging. To represent iNPH brains, another template was created using MRI scans of 22 iNPH patients (14 males, mean age = 71.91±7.06) following the same procedure.

**Fig. 9.**
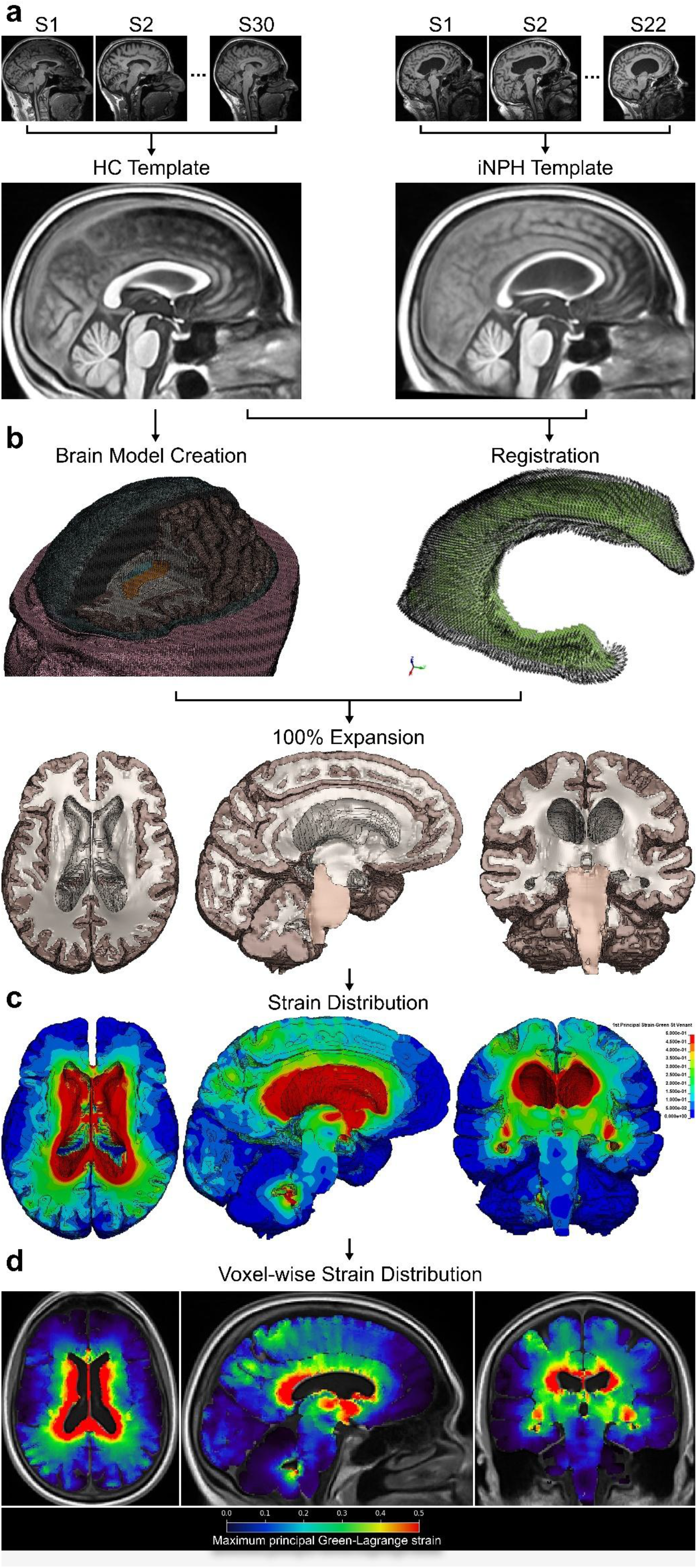
iNPH brain model pipeline. **a)** MRI scans of 30 healthy older adults and 22 iNPH patients were used to generate healthy control (HC) and iNPH templates, respectively, **b)** Anatomically detailed brain finite element model generated using HC template. Registration of HC to iNPH ventricles were used to define a 3D displacement field for lining of ventricles, **c)** Strain distribution show on deformed brain resulted from enlargement of ventricles, **d)** Strain distribution was written in NIfTI MRI format for subsequent analysis, such as extracting strain in white matter tracts.

All individuals participating in this study, including iNPH patients and healthy controls, were scanned at the Imperial College Clinical Imaging Facility using a 3 T Siemens Magnetom Verio Syngo with a 32-channel head coil. A T1-weighted MPRAGE sequence was used with a duration of 5.03 min using the following parameters: “FlipAngle (degrees)” 9, “EchoTime (TR, ms)” 2.98, “RepetitionTime (TR, ms)” 2.300, “InversionTime (TI,ms)” 900, “Bandwidth (Hz/pixel)” 240, “Acquisition matrix” 256 × 240 × 160, “Voxel size (x,y,z; mm)” 1.0, 1,0, 1.0. All iNPH patients had an existing clinical diagnosis and their cases were reviewed a part of this study by a multidisciplinary team of neurologist, neuroradiologists and psychiatrists to confirm their diagnosis.

### Image segmentation

The HC template was segmented using FMRIB and Freesurfer software ^72,73^ for mesh creation. The quality of the segmentation was then checked to ensure the accuracy. This segmentation allows to preserve anatomical features such as sulci and ventricles, which are key for brain biomechanical analyses ^38^. Although the importance of sulci has been demonstrated in a 2D computational model of iNPH ^49^, no prior 3D FE models of iNPH have included sulci. Here, we incorporated sulci in the model to explore their deformation during ventricular expansion. Automatic segmentations of the ventricles can be inaccurate. Hence, all four ventricles were manually segmented in both the HC and iNPH templates using 3D Slicer ^74^. This segmentation included the lower parts of the lateral ventricles and the 3^rd^ and 4^th^ ventricles, which are missing in previous models of iNPH ^44,45,75,76^.

### Brain finite element mesh

The segmented HC template was converted into finite element (FE) mesh using a custom in-house code developed at the HEAD Lab (Fig. 9b) ^41^. This was achieved by creating a hexahedral element at the location of each brain voxel. This approach results in sharp element edges at the surfaces and interfaces, which can cause simulation instability and stress concentration. Hence, a mesh smoothing algorithm was applied on the mesh. The final brain mesh (excluding CSF, ventricles, skin, and skull) was composed of 1,058,711 elements, with an average element size of 0.991 mm, minimum characteristic length of 0.262 mm and maximum characteristic length of 1.374 mm. Mesh quality assessment was conducted using the PyVista library in Python. Results indicated that 98.5% of elements exhibited a good aspect ratio (< 3), 95.2% had an acceptable Jacobian ratio (< 10), and 81.2% maintained an acceptable scaled Jacobian (> 0.5), as illustrated in Fig. 10.

**Fig. 10.**
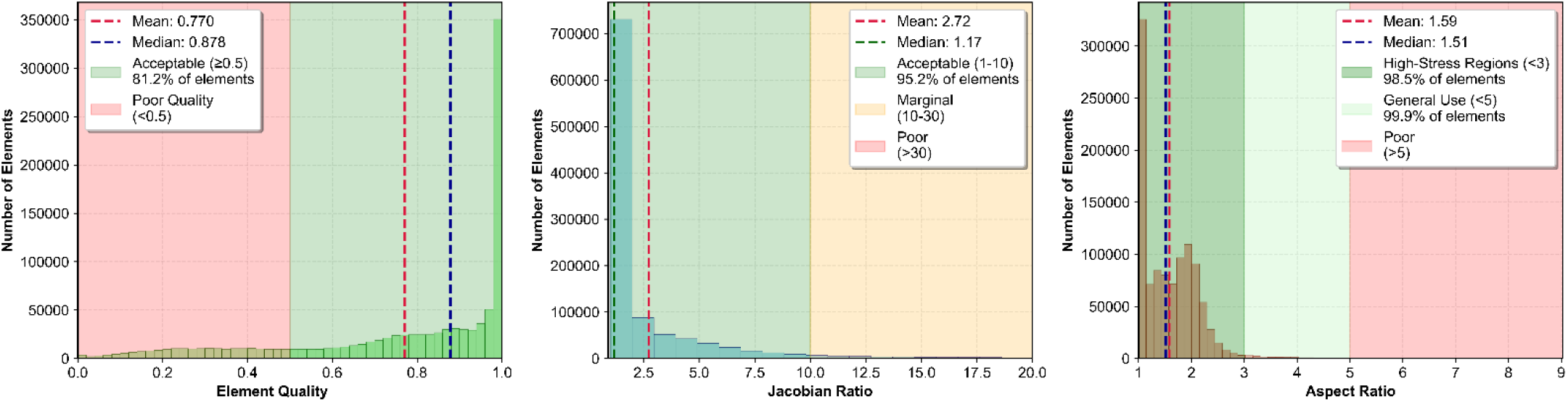
Mesh quality assessment of the brain FE model.

### Other brain structures incorporated in the model

#### Septum Pellucidum

The SP separates the two lateral ventricles, is comprised of both gray and white matter, and is connected to the bottom of the corpus callosum. It typically measures 1–3 mm in thickness ^77^ and has been implicated in neurological disorders ^58^. Despite its significance, no prior iNPH models include a definition of SP. To enhance the biofidelity of our model, a layer of hexahedral elements with an average thickness of 2 mm was added to the mesh using the segmentation refinement pipeline described earlier.

#### Falx

The falx separates the left and right hemisphere and plays a critical role in the brain’s biomechanical behavior ^78,79^. In this study, the falx was modeled as a shell with a thickness of 0.91 mm ^80^. Contact algorithms were implemented to capture interactions between the falx and the corpus callosum as well as between the falx and the left and right brain hemispheres.

#### Pia matter

The pia mater, a thin membrane surrounding the brain, was modeled as a shell with a thickness of 0.2 mm. This layer is essential for maintaining stability at the brain’s outer surface and contributes to overall model accuracy, particularly by its interactions with the inner skull.

### Material model of brain tissue and pia matter

Hyperelastic models, particularly the isotropic Ogden model, have been used to model the stress-strain behaviour of brain tissue under various loading conditions like compression, stretching, and shearing ^81^. Similar to a previous model of iNPH ^43^, we used the Ogden hyperelastic material model, represented by ^82^:

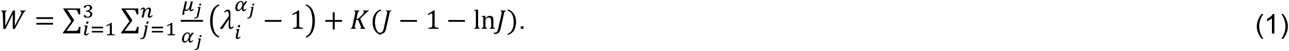

W, 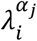, *μ*_*j*_, *α*_*j*_, K and J are potential function, principal stretches, relaxed shear modulus, material coefficient, bulk modulus and volume ratio respectively. The number of terms, *n*, can range from 1 to 8, and for this study, we used *n* = 2. The constants used in this model are provided in detail in Table 1 and Table 2 ^38^.

**Table 1.** Material properties of brain tissue.

| Tissue | Density [$\text{kg}/\text{m}^3$] | $\mu_1$ [Pa] | $\alpha_1$ | $\mu_2$ [Pa] | $\alpha_2$ | Poisson's ratio |
| --- | --- | --- | --- | --- | --- | --- |
| Brain | 1040 | 53.8 | 10.1 | -120.4 | -12.9 | 0.4994 |
| Brain stem | 1040 | 15.8 | 28.1 | -106.8 | -29.5 | 0.4994 |

**Table 2.** Material properties of the head tissues.

| Tissue | Density [$\text{kg}/\text{m}^3$] | $\mu_1$ [kPa] | $\alpha_1$ | Poisson's ratio |
| --- | --- | --- | --- | --- |
| Falx and tentorium | 1130 | 25.5 | 32.9 | 0.45 |
| Pia mater | 1130 | 2.6 | 32.9 | 0.45 |
| | $\tau_i$ [ms] | $\tau_1 = 5$ | $\tau_2 = 44$ | $\tau_3 = 474$ |
| | $G_i$ [kPa] | $G_1 = 32$ | $G_2 = 29$ | $G_3 = 16$ |

We did not include any viscous effects in the material model. This is because we simulated iNPH as a quasistatic problem to account for the long-term progression of iNPH. We modelled the pia mater using a model that combines hyperelastic and viscous effects, similar to previous studies that have used visco-hyperelastic models to simulate quasi-static problems ^83^. This approach prevented numerical instabilities caused by the contact between pia and skull.

### Loading and boundary conditions

We loaded the brain model by expanding the ventricles. In previous FE models of iNPH, ventricular enlargement was simulated by applying pressure on the boundaries of the lateral ventricles. However, this approach cannot accurately reproduce the shape of enlarged ventricles ^22^. Here we apply a three-dimensional displacement field on the lining of ventricles. The displacement field is determined by registering the ventricles of the HC template to the iNPH template. We used two different approaches to achieve most accurate alignment: i) linear and nonlinear image registration (image-based registration), and ii) finite element biomechanical modelling registration (FE-based registration) ^84^.

#### Image-based registration

For the image-based registration, we selected two widely used neuroimaging registration techniques that represent the two principal classes of deformation models: linear (affine) and nonlinear (diffeomorphic) registration. Linear registration was performed using FSL FLIRT, which estimates a global affine transformation based on uniform scaling, rotation and shear between two MRI images. The resulting transformation matrix was applied directly to the HC mesh to generate the approximation of the iNPH ventricle. Nonlinear registration was also performed using the ANTs SyN algorithm, a diffeomorphic approach that models spatially varying, nonuniform deformations. Unlike affine registration, SyN allows each region of the ventricle to deform independently, enabling the algorithm to better accommodate complex geometric changes than FSL FLIRT. This algorithm included an initial affine registration followed by deformable warping. As this method generates a dense voxel-wise displacement field rather than a global transformation matrix, the output could not be applied directly to the HC mesh in the same manner as the linear approach. As a result, the non-linearly registered image of the HC ventricle was directly converted into a mesh to represent the iNPH ventricle instead of extracting any registration matrix or displacement field from the registration process.

#### FE-based registration

FE meshes of the brain were generated from the HC and iNPH templates. The solid elements of the ventricles were extracted from the brain meshes. Then, the shell elements were generated on the surface of ventricles, and the internal solid elements were removed. This process produced paired shell meshes for each ventricle representing the HC and iNPH ventricle.

Next, the shell mesh of the HC ventricle was expanded by applying unit pressure to each element until they reached the fixed iNPH surface. An automatic surface to surface contact algorithm with a soft constraint penalty formulation (scale factor of 0.1) was defined between the two surfaces to prevent interpenetration during expansion simulation. Parameters including material properties and shell thickness were iteratively tuned to maximize numerical stability and achieve optimal geometric registration, rather than to represent physiological tissue properties. Successful registration was achieved using an elastic material model (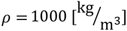, elastic modulus = 10 [kPa], Poisson’s ratio = 0.3) and Belytschko-Tsay membrane element formulation for both meshes. Shell thicknesses were set to 3 mm for the HC and 0.1 mm for the iNPH mesh. The process of FE-based registration is illustrated in Fig. 11.

**Fig. 11.**
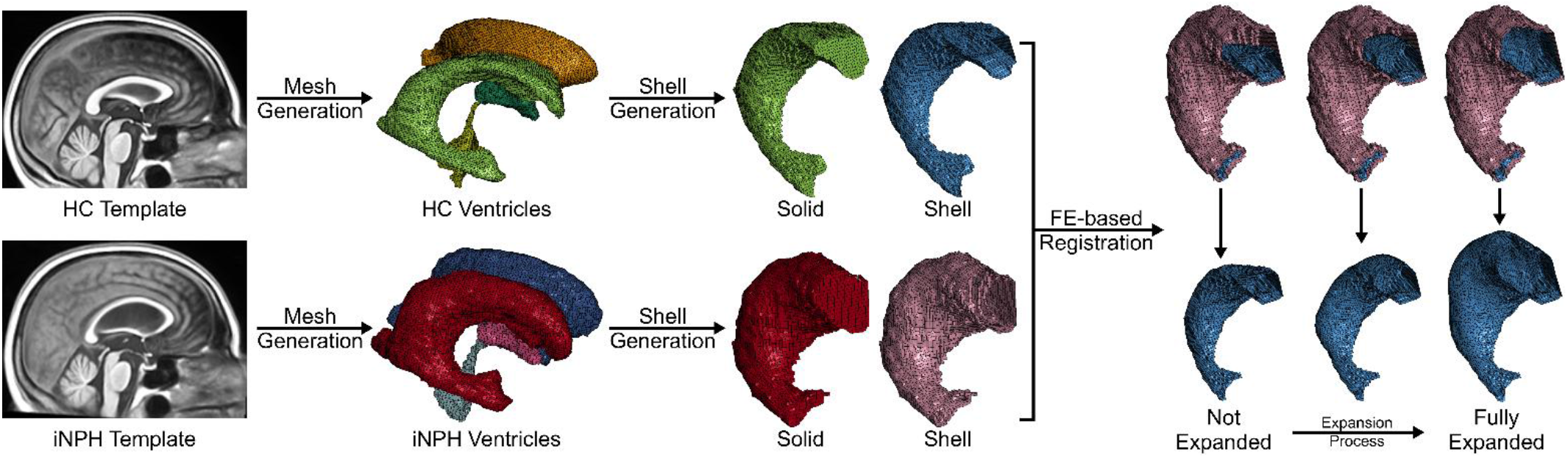
FE-based registration pipeline. iNPH and HC ventricular meshes were generated from templates using an in-house pipeline. Shell surfaces were created for the right ventricles of each group. During registration, the iNPH ventricle was fixed, and the HC ventricle was expanded to conform to the iNPH surface. This registration process was subsequently repeated for other sets of ventricles.

The difference between the position of the HC ventricle nodes before and after expansion was calculated. This resulted in a three-dimensional displacement field for the solid element nodes located on the surface of HC ventricles.

The procedure was repeated independently for both lateral, third and fourth ventricles, resulting in a complete set of nodal displacements describing the transformation from HC to iNPH geometry. The expansion simulations were performed using LS-DYNA, and the displacement vectors were extracted via a custom Python script.

#### Quantifying registration accuracy

Registration accuracy was quantified by using 95^th^ percentile Hausdorff distance (HD95) ^54^. While the standard Hausdorff distance is sensitive to noise, the 95th percentile variant is widely recommended for medical imaging as it mitigates the impact of outliers ^85^. To compute the HD95, the distance from each node on the registered HC ventricle mesh to the nearest point on the reference iNPH ventricle surface was calculated. The HD95 was defined as the distance value below which 95% of these nodal deviations fall. This approach provides a reliable assessment of the spatial correspondence between the registered HC and reference iNPH ventricular geometries, effectively characterizing surface mismatch.

### Controlling kinetic and hourglass energies

We used the nonlinear explicit solver of LS-Dyna to predict brain deformation caused by ventricular enlargement. Explicit solvers can simulate highly nonlinear problems, such as ventricular enlargement, which involves geometric nonlinearity, material nonlinearity and contact. iNPH develops over months to years, but it is not feasible to run LS-Dyna’s explicit solver for such long durations. It is however possible to reduce the loading duration by orders of magnitude as long as the dynamic effects remain negligible. To identify an optimal balance between negligible dynamic effects and loading duration, expansion durations ranging from 10 to 5000 milliseconds were simulated, corresponding to run times of approximately 7 minutes to 60 hours, respectively. Internal and kinetic energy of the brain were monitored. As a rule of thumb, when kinetic energy remained below 5% of the internal energy, we assumed dynamic effects were negligible, preserving the characteristics of iNPH progression (Fig. 12). The results indicated that using 500 milliseconds as the overall expansion time allowed for a faithful representation of the disorder while maintaining computational feasibility.

**Fig. 12.**
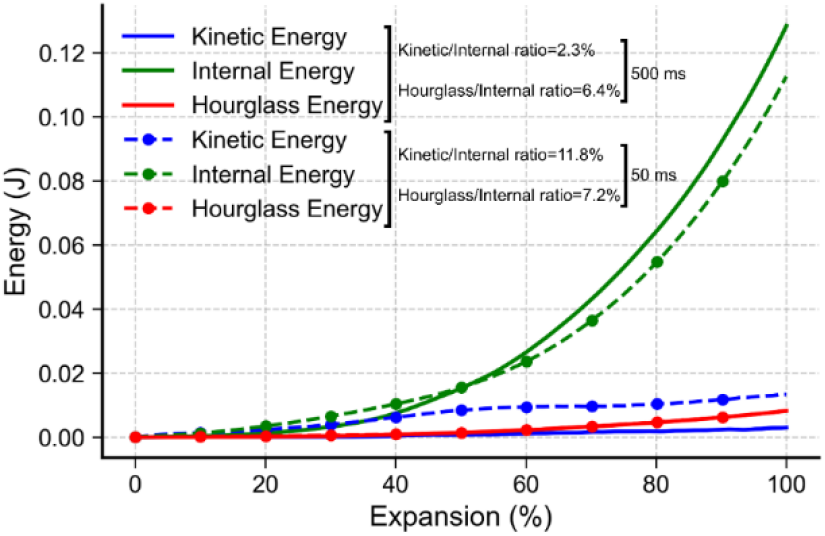
Comparison of internal, kinetic and hourglass energies for two ventricular expansion durations.

To avoid numerical instabilities arising from large shearing deformations and nearly incompressible properties of the brain, we used reduced integration for solid elements of the brain. When reduced order integration is used, spurious deformations, often called hourglass deformation modes, can occur. These deformations should be controlled via using the hourglass control. We applied a stiffness-based hourglass control, with coefficients ranging from 0.01 to 0.5. The hourglass energy, which is an artificial energy produced by the hourglass control, was monitored to ensure it remains less than 10% of the internal energy. When the hourglass coefficient was small, the simulation crashed due to numerical instability (Fig. 13). Increasing the hourglass coefficient improved numerical stability, allowing the simulation to progress further. We selected a coefficient that allowed for the simulation of a fully enlarged ventricle while ensuring the hourglass energy is small enough.

**Fig. 13.**
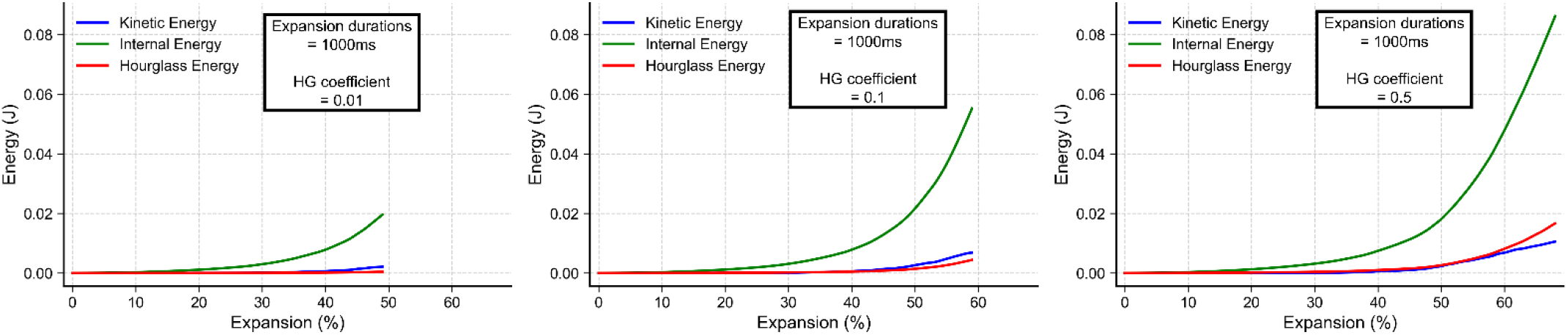
Effect of hourglass settings on simulation accuracy and model stability.

### Tract-based strain distribution

After completion of the ventricular expansion simulation, Green–Lagrange strain was calculated for each element across the brain based on nodal displacements (Fig. 9c). These element-wise strains were converted to voxel-wise strain maps and written in NIfTI MRI format using an in-house pipeline, allowing the biomechanical outputs to be analysed using standard neuroimaging tools (Fig. 9d). To enable tract-based analysis, the JHU ICBM-DTI-81 white matter tract atlas was used to define anatomical regions for sampling the strain values ^70^.

It should be noted that, as the JHU ICBM-DTI-81 white matter tract atlas is provided in MNI152 space (Fig. 14b), it could not be directly applied to the HC template space. To address this, the MNI152 template was nonlinearly registered to the HC template using ANTs. The resulting deformation field was applied to the tract masks, bringing them into HC space (Fig. 14d). These transformed tract masks were then overlaid onto the voxel-wise strain maps, and voxel values within each mask were extracted. For each tract, strain distributions were quantified by computing voxel-wise strain values within the corresponding mask, enabling identification of tract-specific deformation patterns associated with ventricular enlargement.

**Fig. 14.**
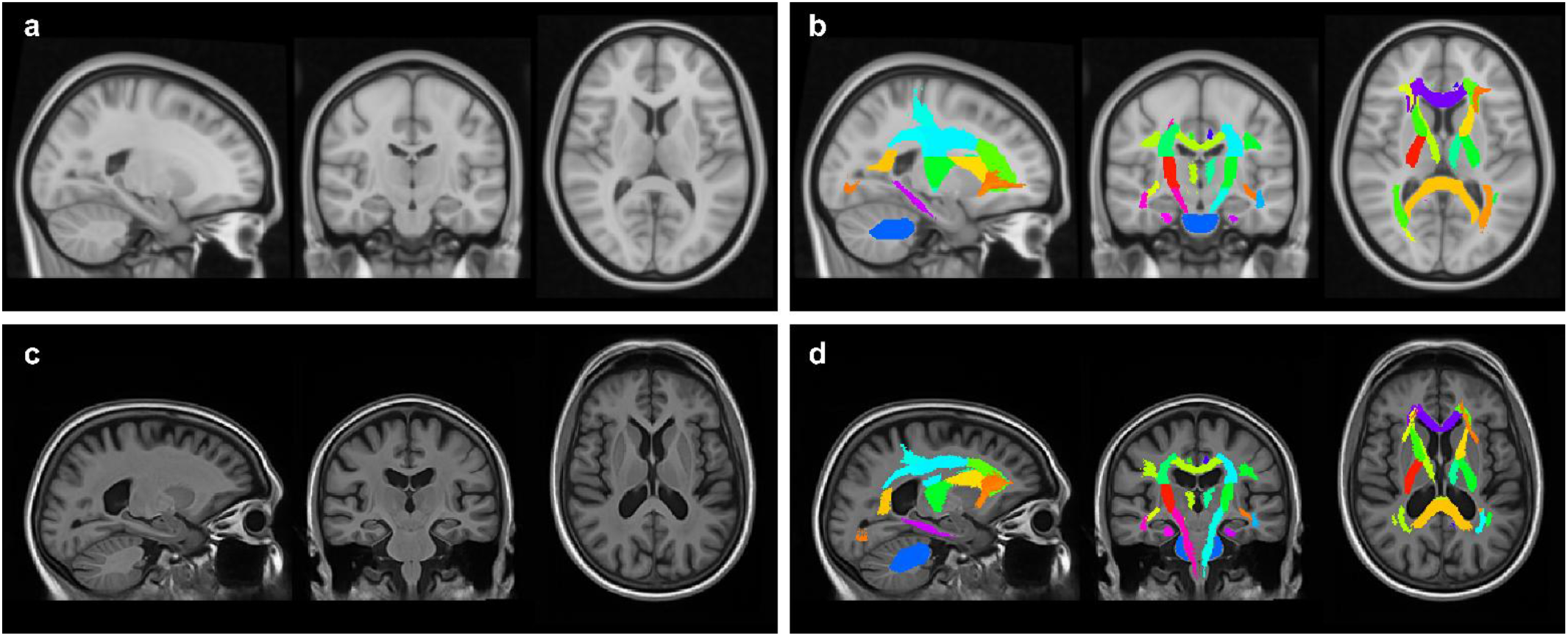
Process of transforming the JHU ICBM-DTI-81 white matter tract atlas into HC space. **a**, MNI152 template. **b**, JHU ICBM-DTI-81 white matter tract atlas in MNI152 space. **c**, HC template. **d**, white matter tract atlas transformed into HC space.

## Ethics and inclusion statement

The acquisition of MRI data for this study was approved by the Health Research Authority’s London-Surrey Borders Research Ethics Committee (19/LO/0102), the London-Central Research Ethics Committee (18/LO/0249), the Camberwell St Giles Research Ethics Committee (17/LO/2066), and the Imperial College Research Ethics Committee (17IC4009). All participants provided digital or written informed consent.

## Data availability

All relevant data supporting the results in this study are available from the corresponding author upon reasonable request and following the completion of a material transfer agreement with Imperial College London.

## Code availability

The finite element simulations were performed using the commercial software LS-DYNA (Version R15, Ansys). Custom code for generating displacement fields and data analysis was implemented in Python (version 3.11.9). All custom code is available from the corresponding author upon reasonable request.

## Acknowledgements

We would like to thank the participants and their families for their invaluable contribution to this research. This work is supported by the UK Dementia Research Institute (DRI) (DRI-CORE2020-CRT) which receives its funding from UK DRI Ltd, funded by the UK Medical Research Council, Alzheimer’s Society and Alzheimer’s Research UK. This work was also supported by the NIHR Imperial College Biomedical Research Centre. D.J.S. has received funding from the Medical Research Council, National Institute of Health Research, Alzheimer’s Society, Football Association, Rugby Football Union, and Premier League Rugby. M.G. acknowledges the support of the Royal Academy of Engineering under the Senior Research Fellowship scheme (RCSRF2324-17-19). V.D. acknowledges the Design Engineering PhD Scholarship, Imperial College London. M.D.G. is employed as research technician at Imperial College London. C.C. is a Consultant Neurologist and Honorary Senior Lecturer at Imperial College Healthcare NHS trust and Imperial College London (MR/W030098/1). M.C.B.D. is an MRC funded Clinical Research Training Fellow (MR/W016095/1). M.A.K. is a Clinical Research Training Fellow supported by the Alzheimer’s Society (608, AS-CTF-22-013). The views expressed in this publication are those of the author and not necessarily those of the NIHR, NHS or the UK Department of Health and Social Care.

## Author contributions

M.G., D.J.S., C.C. and P.A.M. conceived and designed the study. V.D. developed the brain model. V.D. and M.G. designed the method for extracting the new loading conditions. V.D. validated the model, analysed the predictions against radiological data, and conducted the final strain prediction analysis. M.D.G., M.C.B.D., and M.A.K. collected and processed the MRI scans. V.D. and M.D.G. performed registrations. A.G. calculated the radiological scores. M.G., D.J.S., C.C. and P.A.M. supervised the study. V.D., M.G. and M.D.G. wrote the manuscript. All authors reviewed the data and approved the submission.

